# Spatial and single-nucleus transcriptomic profile of a chimpanzee frontal pole

**DOI:** 10.1101/2025.02.16.638510

**Authors:** Gökberk Alagöz, Maggie Wong, Tobias Gräßle, Carsten Jäger, Markus Morawski, Jakob Kolleck, Clyde Francks, Roman M. Wittig, Catherine Crockford, Philipp Gunz, Simon E. Fisher

**Affiliations:** Language and Genetics Department, Max Planck Institute for Psycholinguistics, Nijmegen, The Netherlands; Ecology and Emergence of Zoonotic Diseases, Helmholtz Institute for One Health, Helmholtz Centre for Infection Research, Greifswald, Germany; Department of Neurophysics, Max Planck Institute for Human Cognitive and Brain Sciences, Leipzig, Germany; Paul Flechsig Institute - Centre of Neuropathology and Brain Research, Faculty of Medicine, Universität Leipzig, Leipzig, Germany; Zoological Garden of Landeshauptstadt Saarbrücken, Saarbrücken, Germany; Donders Institute for Brain, Cognition and Behaviour, Radboud University, Nijmegen, The Netherlands; Department of Cognitive Neuroscience, Radboud University Medical Center, Nijmegen, The Netherlands; Ape Social Mind Lab, Institute of Cognitive Science Marc Jeannerod, CNRS, Lyon, France; Department of Human Behavior, Ecology and Culture, Max Planck Institute for Evolutionary Anthropology, Leipzig, Germany; Taï Chimpanzee Project, Centre Suisse de Recherches Scientifiques en Côte d'Ivoire, Abidjan, Ivory Coast; Department of Human Origins, Max Planck Institute for Evolutionary Anthropology, Leipzig, Germany

**Keywords:** chimpanzee, brain, spatial, transcriptomics, genetics, evolution

## Abstract

Chimpanzees, our closest living relatives, share a vast amount of our genetic code, with the majority of differences found in non-coding regions of the genome. Functional and gene regulatory differences drive phenotypic divergence, including the distinctive brain anatomy of humans compared to chimpanzees and other apes. However, little is known about species differences in gene expression, and how they relate to the evolution of neuroanatomy and cognition. This is primarily due to the limited availability of great ape brain samples and challenges in comparative spatial transcriptomic studies. Here, we present the first spatial transcriptomic data from a chimpanzee brain based on *post mortem* tissue from an adult female, who was euthanised due to poor health. We focus on the frontal pole, a brain region that has undergone significant evolutionary changes in size and organisation since the last common ancestor of humans and chimpanzees, and is considered critical for cognitive evolution. We examined the gene expression profiles and cell-type composition of the frontal pole on the left hemisphere, including both neuronal and non-neuronal cell types across cortical layers and white matter. By integrating our spatial transcriptomic data with a publicly available single-nucleus transcriptomic dataset of the chimpanzee dorsolateral prefrontal cortex, we mapped the spatial distribution of 29 chimpanzee brain cell types. This study represents a first step towards characterisation of spatial gene regulatory differences between the brains of non-human great apes and humans.

**Significance statement:** Recent advances in spatial transcriptomics technologies provided important insights into the spatial organisation of gene expression and cell types in human and mouse brains, yet applications in non-model organisms remain limited. Here, we present the first-ever spatial transcriptomics dataset from a chimpanzee brain. We demonstrate i) the applicability of a widely used spatial transcriptomics technique to chimpanzee brain samples freshly frozen in isopentane, and ii) an end-to-end pipeline for generating good-quality spatial transcriptomics data from chimpanzee brains. Our work paves the way for future comparative spatial transcriptomics studies across human and non-human primate brains, and marks a step toward applying spatial omics methods to great ape brains.

## Introduction

Gene expression changes are suggested to be key drivers of phenotypic divergence between closely related species (Wilson et al. 1974; King & Wilson 1975; Enard et al. 2002; Sholtis & Noonan 2010; Reilly & Noonan 2016; Franchini & Pollard 2017), such as humans and chimpanzees. Regulatory changes are often caused by genetic alterations at gene regulatory sequences that impact factors such as chromatin state, transcription factor binding affinity and splicing (Rockman & Kruglyak 2006). These molecular changes can alter the temporal and spatial patterns of gene expression during development and/or in adulthood, and lead to species-specific phenotypes via mechanisms such as acquisition of a new function by a cell type, relocation of an existing cell type and emergence of a completely new cell type (Pollen et al. 2023). In the context of brain evolution, such molecular and cellular changes can lead to microstructural and anatomic changes in white- and grey-matter, eventually driving cognitive and behavioural changes (Rilling et al. 2012; Buckner & Krienen 2013; Schomers et al. 2017). Investigating this long and complex causal chain from regulatory sequence alterations up to neuroanatomic and behavioural changes is quite challenging, but can help to better understand the evolutionary and biological foundations of human neuroanatomy and behaviour.

The advent of RNA-sequencing technologies over the last decade allowed the examination of gene expression at bulk, single-cell/nucleus and spatial levels (Wang et al. 2009; Stark et al. 2019; Rao et al. 2021). Such technological advances fuelled cross-species comparative transcriptomic studies, and led to important discoveries such as a primate-specific striatal interneuron population (Krienen et al. 2020) and genes expressed in primary motor cortex with human-specific co-expression patterns (Suresh et al. 2023). Despite such important insights, differences in transcriptional and developmental programmes shaping human and non-human great ape neuroanatomy are largely unknown. Drawing on prominent views on human brain evolution and recent findings from molecular evolution studies, Pollen et al. (2023) suggested that shedding new light on the spatiotemporal gene expression programmes involved in human and non-human great ape neurodevelopment is crucial to better understand how the human-specific aspects of our brains have evolved.

The recent availability of spatial transcriptomics methods enabled the investigation of spatial gene expression patterns, as well as the localisations of and interactions between different cell types (Vandereyken et al. 2023; Ma & Zhou 2024; Pham et al. 2023). While spatial transcriptomics data from various cortical and sub-cortical human brain regions have accumulated over the last few years (Chen et al. 2020; Maynard et al. 2021; Wong and Sha et al. 2024; Huuki-Myers et al. 2024), such insights from non-human great ape brains are completely missing due to the scarcity of good-quality great ape brain samples.

Here, we present the first spatial transcriptomics data from a non-human great ape brain, alongside a comprehensive pipeline – from necropsy to data generation and analysis – for high-quality, fresh-frozen chimpanzee brain spatial transcriptomics. The brain tissue was opportunistically obtained from an adult female chimpanzee at the Saarbrücken Zoo (Germany), which had to be euthanized due to a poor medical prognosis (unrelated to our study). We focused our efforts on the left hemisphere frontal pole, which is a region known to be substantially different in size and organisation between humans and chimpanzees (Semendeferi et al. 2001, Semendeferi et al. 2011), and is suggested to be involved in various aspects of cognition such as self-awareness, decision-making and metacognition (Mansouri et al. 2017; Bouret et al. 2024). By combining our fresh-frozen brain sample preparation method with 10x Genomics Visium, a well-established sequencing-based spatial transcriptomics method (Ståhl et al. 2016; Salmén et al. 2018), and a set of analytic tools, we were able to generate good quality chimpanzee brain spatial transcriptomics data. Finally, we integrated our Visium data with a publicly available single-nucleus transcriptomics dataset of chimpanzee dorsolateral prefrontal cortex using a Bayesian model implemented in cell2location (Kleshchevnikov et al. 2022), and annotated cell type abundances on chimpanzee frontal pole. Our study establishes a comprehensive framework for investigating spatial gene regulatory profiles of great ape brains, and paves the way for future great ape brain comparative spatial transcriptomics studies.

## Results

### Histological and transcriptomic quality control of chimpanzee brain tissue sections and comparison with publicly available human data

We first performed a set of qualitative and quantitative controls to ensure the suitability of our sample for the Visium protocol. We measured the integrity and concentration of RNA isolated from two tissue samples obtained from two tissue blocks in the left frontal pole, and observed that both RNA integrity and concentration measures were sufficient to proceed with spatial transcriptomics data generation (Table S1). We then performed cryosectioning on two adjacent left hemisphere frontal pole tissue blocks, and obtained two pairs of 10-µm thick consecutive tissue sections, one pair from each block (Fig. 1A, Fig. S1). We placed these tissue sections on a Visium spatial gene expression slide and performed haematoxylin and eosin (H&E) staining to characterise the anatomical morphology, revealing roughly marked white matter-grey matter borders, blood vessels and gyrification patterns, as expected (Fig. 1B).

**Figure 1:**
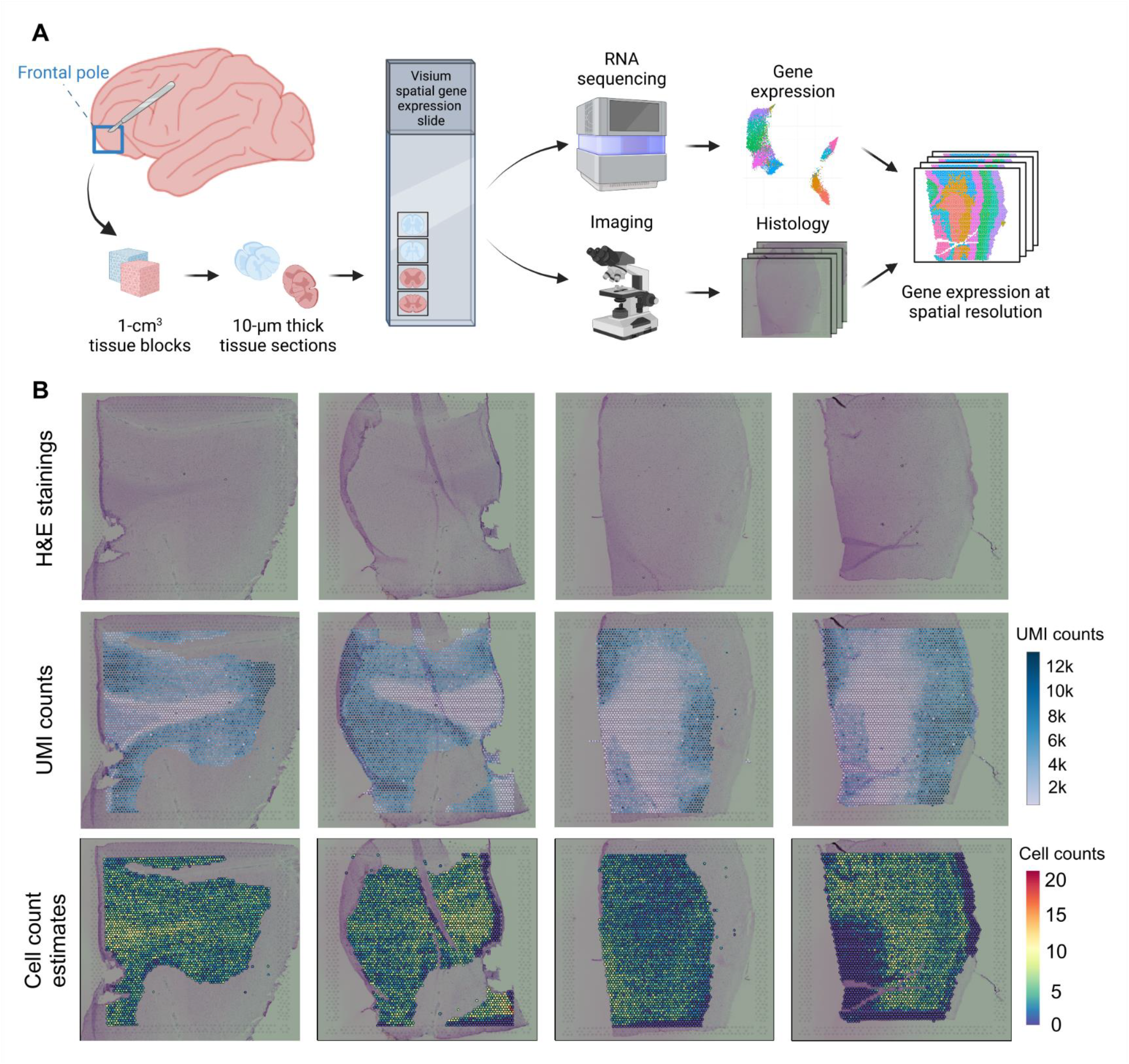
Spatial transcriptomics workflow and an overview of the chimpanzee frontal pole tissue sections. **(A)** Schematic shows an overview of our spatial transcriptomics data generation and analysis pipeline. **(B)** H&E staining of all sections that are investigated in this study, depicting the overall tissue morphology (haematoxylin stains nuclei with purple, eosin stains extracellular matrix and cytoplasm with pink). Unique molecular index counts and cell count estimates are depicted as spot-plots.

We then performed spatial transcriptomic profiling (median sequencing depth=283×10^6^, sequencing depth range=221×10^6^-316×10^6^), and measured gene expression of 23,012 genes in 13,399 spatially barcoded Visium spots across four sections (Table S2). Importantly, our analysis yielded a mean of 5,002 unique molecular identifiers (UMI) and 1,879 genes per Visium spot, which is close to that of human brain-derived spatial transcriptomics data available in literature (Fig. 1B, Table S3) (Chen et al. 2020; Maynard et al. 2021; Wong and Sha et al. 2024; Huuki-Myers et al. 2024). We observed that the UMI count distribution patterns differ between white and grey matter, a phenomenon also reported in a prior human brain spatial transcriptomics study (Wong and Sha et al. 2024) which is potentially due to the lower RNA content in axons compared to neuronal cell bodies.

To further investigate the overall morphology of the chimpanzee frontal pole, we quantified the nucleus density in our tissue sections and compared our findings to a publicly available human dorsolateral prefrontal cortex dataset (Maynard et al. 2021). Our image analysis, which leveraged the high-resolution H&E staining images, identified 2.7 nuclei per Visium spot on average across all spots from four tissue sections (Table S3). We also found that 9.5% of all spots harboured a single nucleus, whereas 10.3% did not contain any nuclei (Fig. S2). These measures are similar to the human brain-derived nucleus count estimates in Maynard et al. (average nuclei per spot=3.3, percentage of spots with one nucleus=15.0, percentage of spots with zero nucleus=9.7) (Maynard et al. 2021).

Before conducting spatial transcriptomics data analyses, we performed immunohistochemical protein staining to gain an initial overview of the cytoarchitecture in our chimpanzee frontal pole samples. We stained eight tissue sections, which were obtained from the same two tissue blocks used for spatial transcriptomics. To reveal the white matter-grey matter boundary and to highlight deep- and outer-layer neurons, we stained for canonical protein markers that are known to be abundant at certain layers of the human cortex (Zeidán-Chuliá et al. 2016; Maynard et al. 2021), including a pan-neuronal marker (NeuN), two deep layer neuronal markers (CTIP2 and TBR1), an upper layer marker (BRN2), and a white matter marker (MBP). Layer-specific localisations of canonical markers in human brain studies were rather ambiguous in our staining images (Fig. S3). However, the white matter-grey matter distinction that we observed in H&E images and UMI count distributions overlapped with the spatial patterns revealed especially by anti-NeuN, -TBR1 and -BRN2 staining.

### Spatial variation in gene expression reflects grey-white matter distinction and the laminar structure in grey matter

After performing quality control to ensure data quality was comparable to that of publicly available human brain-derived spatial transcriptomics data, we integrated and normalised the spatial transcriptome data from our tissue sections by using Harmony (Korsunsky et al. 2019) to correct for batch effects (Fig. S4). We applied further filtering on genes and Visium spots (as described in Methods), and removed spots located under folded tissue sections, which yielded 12,614 spots in total. We then identified 12 gene expression clusters of Visium spots from across all tissue sections by using a Markov Chain-Monte Carlo clustering algorithm provided in BayesSpace (Zhao et al. 2021) (Fig. 2A, Fig. S5). To investigate the spatial distribution of these transcriptomically-defined clusters, we mapped the cluster identity of each Visium spot back to its corresponding histology image. We observed that the 12 gene expression clusters mainly reflect i) transcriptomic differences between white and grey matter, and ii) cellular and gene expression variability across layer-like laminar structures within the grey matter of the chimpanzee frontal pole (Fig. 2B). This transcriptomically-defined laminar spatial organisation, including the specific order of gene expression clusters across the axis spanning from the white matter towards the outer layers of grey matter, was consistent among all four sections. Thus, these clusters appear to reliably capture the distinct molecular characteristics of the various cortical layers.

**Figure 2:**
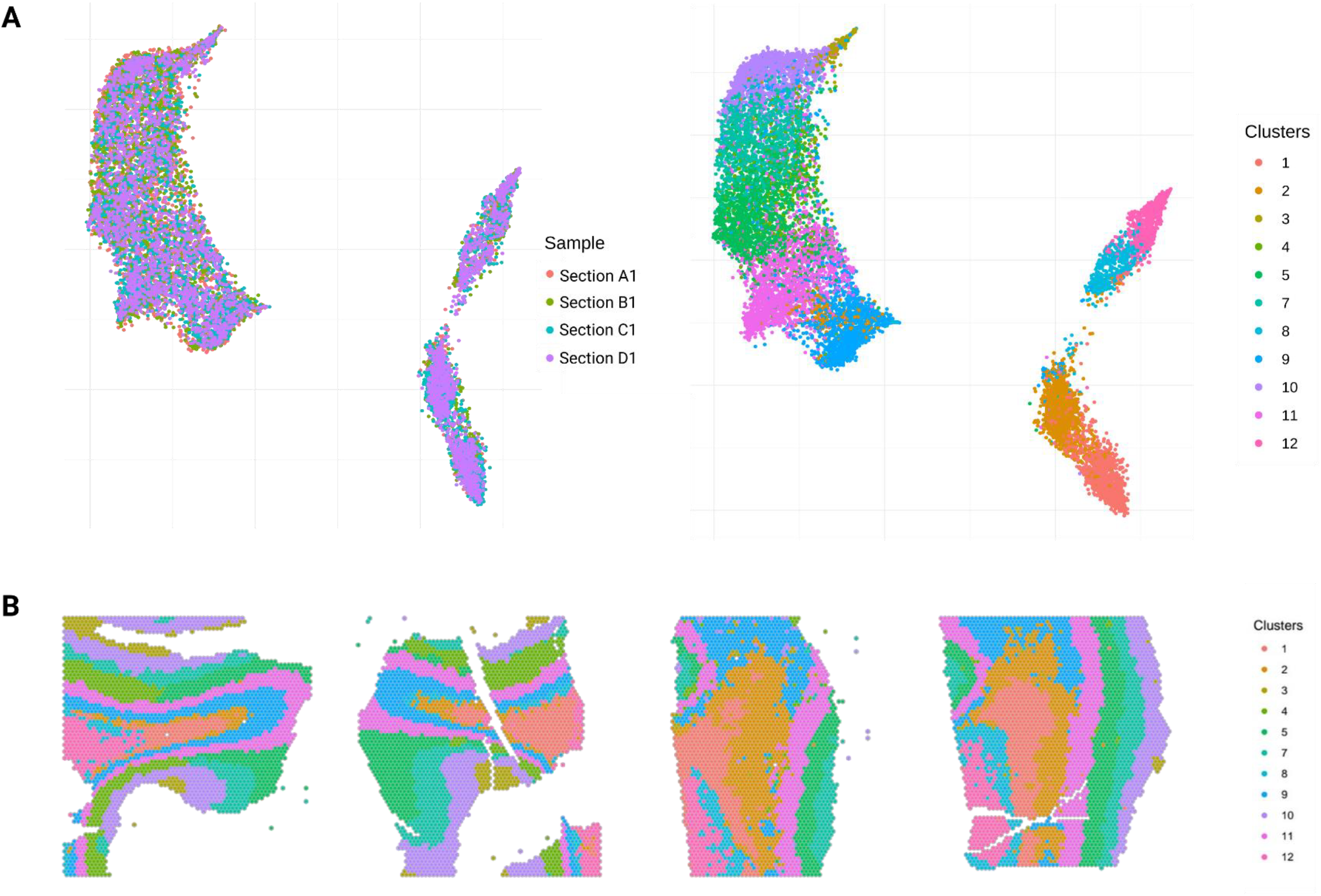
UMAP and spatial representations of gene expression clusters. **(A)** Unsupervised clustering of Visium spots based on their gene expression profiles after integration of data from four sections. Colours indicate which tissue section each spot belongs to (left) and 12 gene expression clusters identified by the BayesSpace (right). **(B)** Spatial map of gene expression clusters on histology images. Colours correspond to the same 12 gene expression clusters with the UMAP on top right.

### Integration with a publicly available single-nucleus RNA-sequencing dataset and cell type deconvolution analyses reveal cell subclass organisation in the chimpanzee frontal pole

To investigate the spatial organisation of various brain cell types in the chimpanzee frontal pole, we deconvoluted our spot-level data to cellular level. To do so, we integrated the Visium spatial transcriptomics data with a publicly available single-nucleus RNA-sequencing (snRNA-seq) dataset (Ma et al. 2022) derived from dorsolateral prefrontal cortex tissue samples of four adult chimpanzees (158,099 nuclei). This reference dataset contains annotations of 29 neuronal, glial, and non-neuronal cell subclasses (Table S4) based on gene expression profiles, providing a high-resolution cellular map of the chimpanzee dorsolateral prefrontal cortex, which is a large cortical area located adjacent to the frontal pole (Jung et al. 2022).

We first quantified RNA and cell abundances to estimate RNA detection sensitivity per-spot for all sections using cell2location (Kleshchevnikov et al. 2022) (Fig 3A, Fig. S6-7). Spatial variation in RNA detection sensitivity estimates overlapped with the spatial distribution pattern of UMI counts, which indicates that low RNA counts in white matter can hinder cell2location’s RNA detection sensitivity. We then generated a cell subclass-specific gene expression signature reference by training a Bayesian model on the single-nucleus gene expression data from Ma et al. (Ma et al. 2022) using cell2location (Kleshchevnikov et al. 2022). We integrated these reference gene expression signatures with our spatial transcriptomics data, and trained an RNA detection sensitivity-aware model to estimate per-spot cell subclass abundances for 29 cell subclasses (Fig. S8). Overlaying the abundance of four neuronal subclasses with layer identity (layer 2-3 intratelencephalic, layer 3-5 intratelencephalic type 1, layer 3-5 intratelencephalic type 3, and layer 6 corticothalamic neurons), as well as oligodendrocytes and astrocytes, further highlighted the laminar structure of the chimpanzee grey matter, and the distinction between grey and white matter. The localisations of four selected neuronal subclasses converged with the spatial organisation of 12 gene expression clusters identified by unsupervised clustering (Fig. 3B), which suggests that these clusters also have layer identities potentially with unique cellular compositions. The spatial order of the four neuronal subclasses with cortical layer identities, as identified by Ma et al. (Ma et al. 2022), match their expected anatomical order along the axis from the outer zone of grey matter to white matter, again supporting the convergence of reference data-based cell subclass mapping and unsupervised clustering analysis results.

**Figure 3:**
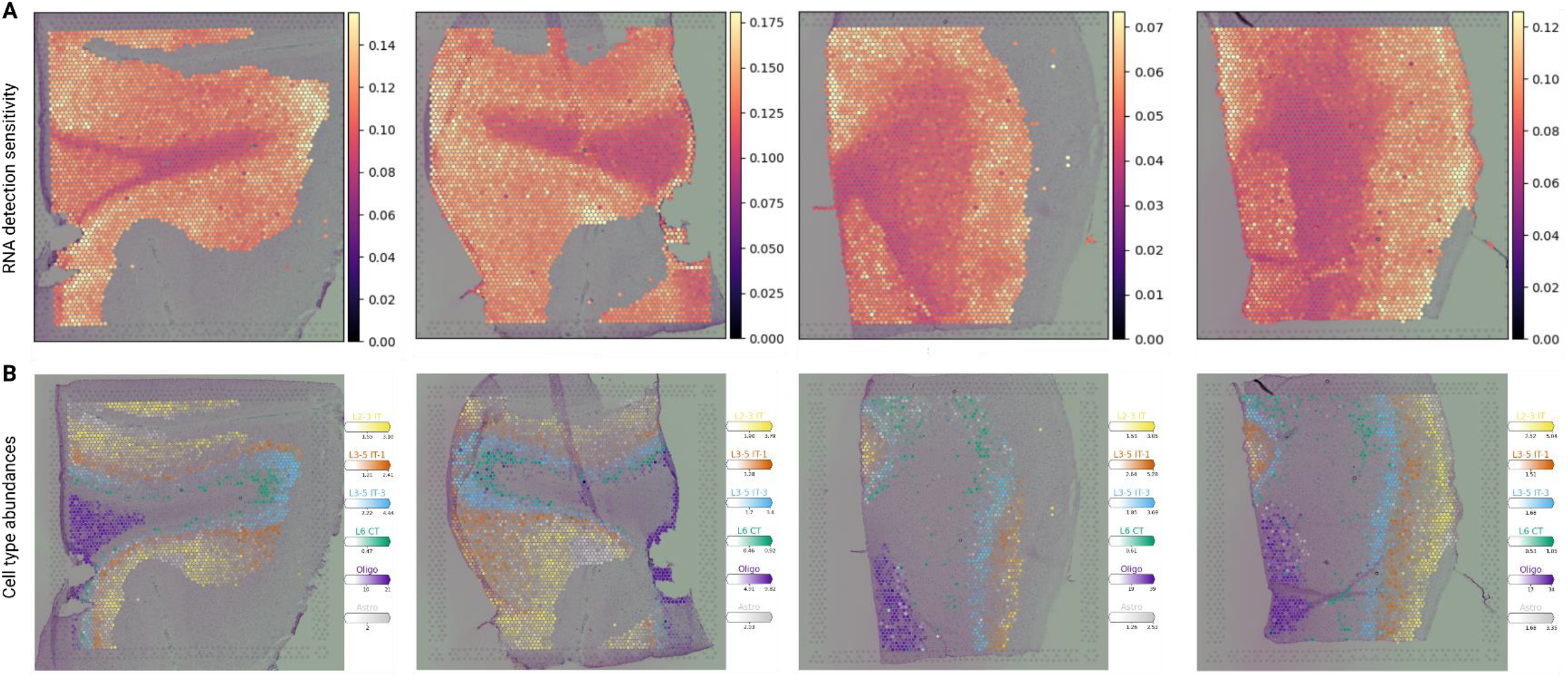
RNA detection sensitivity and cell subclass abundance estimates by cell2location. **(A)** Per-spot RNA detection sensitivity measures for each section. **(B)** Cell subclass abundances for six selected subclasses [L2-3 IT (yellow): Layer 2-3 intratelencephalic; L3-5 IT-1 (orange): Layer 3-5 intratelencephalic Type 1; L3-5 IT-3 (blue): Layer 3-5 intratelencephalic Type 3; L6 CT (green): Layer 6 corticothalamic; Oligo: Oligodendrocytes (purple); Astro: Astrocytes (grey)].

## Discussion

Here, we have established and validated in pilot experiments a workflow to study spatial gene expression in great ape brains, covering steps from necropsy, tissue extraction and freezing, through to spatial transcriptomics data generation and analysis. We successfully generated high-quality spatial transcriptomics data from a chimpanzee brain using the Visium platform, with comparable RNA quality and quantity measures to that of publicly available human brain-derived data. We have applied a set of integrative omics tools to our dataset, and successfully profiled the spatial gene expression patterns and cell type composition in the chimpanzee frontal pole.

We compared a number of standard spatial transcriptomic data quality and quantity measures between our chimpanzee brain-derived data and three publicly available human brain-derived spatial transcriptomics datasets (Chen et al. 2020; Maynard et al. 2021; Wong and Sha et al. 2024; Huuki-Myers et al. 2024). These comparisons provided a key checkpoint before proceeding with data analysis, as the experimental and analytic protocols for spatial sequencing technologies were not tested on non-human great ape brains prior to this study. We observed that the average number of unique RNA molecules, genes and cells per Visium spot in chimpanzee frontal pole and human cortex tissue sections are similar, suggesting that spatial transcriptomics data from these two species can be analysed in a comparative design. Our findings support previous observations (Wong and Sha et al. 2024) showing that the number of unique RNA molecules per-spot in white matter is lower compared to grey matter, which might be related to the lower RNA content in the axons compared to cell bodies. Yet, we note that the sample size of our study does not have sufficient statistical power to reach generalisable biological conclusions on the molecular characteristics of the chimpanzee frontal pole. Nonetheless, obtaining similar data quality and quantity measures with prior human brain spatial transcriptomics studies enabled us to proceed with spatial gene expression data analysis. Previously, such comparative spatial transcriptomics studies were performed between human and mouse samples, revealing distinct cell types and species-specific organisation of gene expression (Beauchamp et al. 2022). Our study paves the way for similar comparative studies across humans and non-human great apes, which may provide important insights into species-specific aspects of our brains.

Our gene expression data-driven clustering analysis identified spots with distinct transcriptomic profiles. The main axes of gene expression variation were driven by the transcriptomic and cell type composition differences between white and grey matter, and across cortical layers. Despite the limited sample size of our study, this observation is in line with various human brain spatial transcriptomics studies, indicating the overall spatial patterns of gene expression can be recapitulated even in small datasets. Similarly, our observation that the gene expression clusters are organised in a laminar pattern is in line with the well-known six layered structure of the mammalian cortex (Molnár et al. 2014). To support our findings from unsupervised clustering, we performed single-nucleus data integration and cell type abundance estimation analysis, which replicated this laminar distribution and showed that the gene expression clusters overlap with cell subclasses with cortical layer identity in the expected layer order (Akula et al. 2023). Our results from the unsupervised clustering and the reference data-based cell type mapping analyses converged, validated the histologically identified laminar structure of the chimpanzee cortex (Balaram et al. 2014; Amiez et al. 2021), and provided a first glance into the fine-grained cell type organisation in the chimpanzee cortex.

The main limitation of the present study is that tissue sampling was restricted to the left hemisphere frontal pole of a single female chimpanzee brain. Thus, even though we were able to observe the laminar organisation of gene expression in the adult chimpanzee frontal pole, we refrain from drawing cortex-general, species-general or comparative conclusions. In addition, the absence of a chimpanzee reference transcriptome, and the incomplete annotation of the chimpanzee genome may introduce a bias to quantify the expression of genes that are well-annotated. Ongoing efforts to generate high quality and high coverage primate genome assemblies (Kuderna et al. 2023; Housman & Tung 2023) hold great promise for the availability of better annotated great ape genomes in the near future that will help to avoid this bias in future studies. Secondly, the relatively long *post mortem* interval prior to brain extraction (27 hours) and freezing may lead to RNA degradation. To circumvent this potential limitation, we cooled down the body immediately after euthanasia, and managed to retrieve non-degraded RNA from the brain samples. Finally, we were not able to recover sufficient UMI and gene counts from some Visium spots, potentially due to technical reasons including uneven freezing pace of different parts of the tissue sections and condensation formed due to fluctuations or suboptimal temperature fluctuations within the cryostat when transferring cryosections onto Visium slides. We recommend using a cryostat with a stable chamber, holder and blade temperature to overcome similar issues.

In summary, our study represents a promising first step towards the spatial transcriptomic profiling of non-human great ape brains. We believe that the rapid enhancement of the spatial biology technologies (Marx 2021) and efforts like ours to apply these methods to non-human great apes pave the way for the generation of comprehensive great ape brain cell atlases. Combined with the ongoing efforts of human brain cell atlases (Wang Daifeng et al. 2018; Huuki-Myers et al. 2024), future comparative spatial research on human and non-human great ape brains can shed light on the spatiotemporal characteristics of gene expression, which drive human-specific aspects of neurobiology and cognition.

## Materials and Methods

### Necropsy and sample preparation

For this study, we collected a 53-year-old female captive chimpanzee (*Pan troglodytes troglodytes*) brain in July 2021 at the Saarbrücken Zoo in Germany, following her euthanisation due to poor medical prognosis caused by a right abdominal cavity tumor. The body was cooled at 3-5 °C, and the brain was extracted 27 hours *post mortem*. We first cut apart the cerebellum and the cerebrum. The left and right hemispheres were then separated by cutting the corpus callosum. Left and right hemispheres were sliced coronally into ∼1-cm thick slices, which were freshly frozen in isopentane (−80 °C). Then, half of the rostral-most slice of the left hemispheric cerebrum (i.e. the frontal pole) was cut into five ∼1 cm^3^ tissue blocks on dry ice. After visually inspecting these tissue blocks’ suitability (e.g. having large and flat surface) for cryosectioning and Visium spatial transcriptomics protocols, we selected two tissue blocks for RNA quality and quantity control measurements, and another two blocks for performing spatial transcriptomics (see Fig. S1 for details) and immunohistochemistry.

### Sample processing and Visium data generation

We cryosectioned (CM1900, Leica) the selected frozen tissue blocks at a chamber and blade temperature of -20 °C, and a holder platform temperature of -12 °C. We took two 10-µm thick consecutive sections from each block, obtaining four tissue sections in total. Sections were placed on a pre-chilled (down to -20 °C) Visium Spatial Gene Expression Slide (Visium Spatial Gene Expression Slide & Reagent Kit, 10x Genomics, kit no. 1000184, Visium slide no. V13J17-280), and immediately attached sections to the slide by warming the back of the slide. Following the Methanol Fixation, H&E Staining and Imaging for Visium Spatial Protocols (catalogue no. CG000160 Rev C), we immersed and incubated our Visium slide in pre-chilled methanol (−20 °C) for 30 minutes to fixate the tissue sections, after which we carried out H&E staining and tissue imaging. We took brightfield images of H&E stained sections using an AxioScan Z1 slide scanner (Zeiss) with a Hitachi HV-F292SCL camera, a Plan-Apochromat 20x/0.8 M27 objective and Zeiss Zen v.2.6 (Blue Edition) software.

We then proceeded with permeabilisation of the tissue sections (for 18 minutes) and reverse transcription, followed by second strand synthesis and denaturation following the Visium Spatial Gene Expression User Guide (catalogue no. CG000239 Rev F, 10x Genomics). We performed qPCR to determine the number of cDNA amplification cycles required to reach ∼25% of the peak fluorescence value, and then ran 16 cycles of cDNA amplification. Finally, we performed library preparation using the BioRad T100 Thermal Cycler for PCR reactions following the Spatial Gene Expression Library Construction section of the Visium Spatial Gene Expression User Guide (catalogue no. CG000239 Rev F, 10x Genomics). The libraries were sequenced on an Illumina NovaSeq 6000 System with 150 bp paired-end reads using the standard sequencing protocol for Visium fresh-frozen libraries (Read 1: 28 cycles, i7 index: 10 cycles, i5 index: 10 cycles, Read 2: 91 cycles). The four libraries were sequenced to a median depth of 283×10^6^ reads (range: 221×10^6^-316×10^6^).

### Histology image analysis and nucleus segmentation

Following the methodology provided by Maynard et al. (2021) (also see https://www.mathworks.com/help/images/color-based-segmentation-using-k-means-clustering.html), we applied K-means colour-based nucleus segmentation to the high-resolution H&E staining images. We first converted each image from RGB to CIELAB colour-space by using the rgb2lab function in Matlab (vR2019a). CIELAB colour-space quantifies luminosity measures (from black to white), as well as two chromacity spectrum measures (along red-green and blue-yellow axes). We extracted two chromacity layers from each image, which allowed us to visually inspect and identify the number of different colours in each image. We then used the imsegkmeans function to generate binary masks for each manually observed colour in the histology images, which allowed us to obtain a binary mask per section tagging all nuclei which were dyed in purplish blue by haematoxylin. Following the nuclei number estimation per Visium spot protocol implemented by Maynard et al., we integrated these binary masks with individual spot coordinates that we obtained from data processing by SpaceRanger. By overlaying the binary masks with Visium spot coordinates, we estimated the number of nuclei that fell in each spot and generated a summary table including per-spot estimates.

### Visium spatial transcriptomics data processing

We integrated raw sequencing data (FASTQ files) and the low-resolution H&E images to perform read alignment against the chimpanzee reference genome Clint_PTRv2 (a.k.a. pan_tro6; accession number: GCF_002880755.1). We rescaled the H&E images to 2000x2000 pixels by using ImageJ (v1.53). We then performed a standard QC of the sequencing data using default parameters in SpaceRanger (v1.2.2) (see Table S1 for data quality control measures). The resulting count matrices (feature-barcode matrices), which contain UMI counts of each gene-spot pair, were then used for data analysis.

Following the analysis pipeline used by Maynard et al. (Collado-Torres et al. 2020), we imported mapped and preprocessed Visium data and the matching low-resolution H&E image for each sample into R using SummarizedExperiment (Martin Morgan 2017). We obtained a large SingleCellExperiment object which includes data from 13,399 Visium spots across four sections, with a mean of 5,002 UMIs and 1,879 genes per spot. First, we performed spot-level quality controls of UMI count, gene count and percentage of mitochondrial reads by using perCellQCMetrics and quickPerCellQC functions from the Scran package (v1.18.7). We did not filter out spots based on these measures, as the UMI and gene count variance across spots largely reflected biological differences between grey and white matter. We then removed the spots located under folded parts of tissue sections, resulting in high-quality data from 12,614 spots across four sections. Second, following the data filtering protocol from Wong and Sha et al. (Wong and Sha et al. 2024), we performed gene filtering by removing the genes that are expressed in less than 0.01% of all spots, and all mitochondrial genes. A total of 23,012 genes passed these filtering steps.

### Visium spatial transcriptomics data analysis

In order to prepare raw data for unsupervised gene expression clustering, we first computed the log-normalised gene expression counts per spot and estimated the first 50 principal components based on gene expression variation among the top 2,000 highly variable genes using the spatialPreprocess function from the BayesSpace R package (Zhao et al. 2021). We then performed a principal component analysis to estimate the top 50 axes of gene expression variation across Visium spots from four sections using the runUMAP function of the scater R package (McCarthy et al. 2016). We first used these top 50 principal components to perform Uniform Manifold Approximation and Projection (UMAP) dimensionality reduction and explored data variation axes prior to data harmonisation. We then integrated principal component analysis results with the main SingleCellExperiment object, and corrected for technical variation across spatial transcriptomics libraries (i.e. batch effects) using the runHarmony function from the Harmony R package (Korsunsky et al. 2019), which took the log-transformed gene expression count data combined with the top 50 principal components. By combining this enhanced data object with user-specified batch or sample information, Harmony corrected for technical variation in our data while preserving transcriptomic variation. We generated a new UMAP after data harmonisation to ensure that batch effects did not impact main axes of variation in our data.

Next, we selected the number of gene expression clusters to use in our clustering analysis by using the qTune function of BayesSpace. We estimated average pseudo-log-likelihoods for a set of cluster numbers (q=2 to 20), plotted log-likelihoods as a function of q and chose the q value around the elbow of this plot (q=12) as the number of gene expression clusters. We then performed clustering analysis for all spots across four sections using the spatialCluster function of BayesSpace (q=12, total number of iterations=20,000, burn-in iterations=1,000, seed number=149). We projected these data-driven gene expression clusters of Visium spots onto H&E images and observed the spatial distribution of 12 gene expression clusters across four sections using the clusterPlot function of BayesSpace.

### Single-nucleus RNA-sequencing data-supervised cell type deconvolution analysis

We used a Bayesian model named cell2location (Kleshchevnikov et al. 2022) to integrate our Visium spatial transcriptomics data with a publicly available chimpanzee dorsolateral frontal cortex-derived single nucleus-RNA sequencing dataset (Ma et al. 2022). The dataset from Ma et al. contains transcriptomic data from 29 neuronal, glial and non-neuronal cell subclasses (see http://resources.sestanlab.org/PFC/). Following the cell2location tutorial for Visium spatial transcriptomics data analysis (see https://cell2location.readthedocs.io/en/latest/notebooks/cell2location_tutorial.html), we imported mapped and pre-processed Visium data, and the matching low-resolution H&E images for each sample, into Python (v3.9) by using the read_visium function from the scanpy package (Wolf et al. 2018). We filtered out mitochondrial genes from our Visium data.

Next, we downloaded processed single-nucleus RNA-sequencing data from the Ma et al. study (see http://resources.sestanlab.org/PFC/), imported the raw count matrix to Python, and removed non-chimpanzee samples from the data. We applied a permissive gene selection in this dataset by excluding genes with less than 1.1 mean expression among all spots and genes that are not expressed at least in 0.03% of all spots, which yielded 13,279 genes. Then, we estimated reference cell subclass signatures using cell2location, which follows a two-step procedure: i) negative binomial regression to combine and harmonise single-nucleus RNA-sequencing data across batches/samples, and ii) computation of average gene expression levels per gene for each cluster by training a model (training epochs=250). We performed standard quality control steps to check the accuracy of this model by checking the evidence lower bound (ELBO) loss history and reconstruction accuracy, and then proceeded with integrating the reference data set with our Visium data.

We performed cell type deconvolution analysis for each Visium section independently by using the default cell2location model to estimate cell type abundances in each Visium spot. We informed this model with the average per-spot cell number estimate that we obtained from the aforementioned high-resolution image analysis. Once the model was trained (training epochs=30,000), we performed a quality control to check the ELBO loss history and reconstruction accuracy measures (Fig. S6), and spatially visualised cell abundances for 29 cell subclasses by using the 5% quantile of the posterior distribution, which represents cell abundances in which the model has high confidence.

### Immunohistochemistry

Cryosections (10-µm thick) adjacent to those used for Visium were fixated in 4% PFA at room temperature for 10 minutes, followed by 1X PBS washes (3 x 5 minutes). Then, we applied antigen retrieval by incubating sections in pre-warmed citrate buffer (pH 6.0) (Merck, C9999) at 90 °C for 10 minutes in a water-bath, which were then cooled down to room temperature for 30 minutes. Sections were washed again with 1X PBS (3 x 5 minutes). After washing, we drew hydrophobic barriers between sections using an immunostaining pap pen (Kisker Biotech, MKP-1) to prevent cross-binding of different antibody combinations. Sections were then incubated in blocking and permeabilisation buffer containing 5% donkey serum (Sigma, S30-100ML) and 0.25% Triton (Sigma Aldrich, X-100) in PBS at room temperature for one hour.

Primary antibodies: NeuN (Millipore, MAB377, 1:2000), BRN2 (Cell Sig., 12137S, 1:500), CTIP2 (Abcam, ab18465, 1:500), MBP (Sigma, NE1019-100UL, 1:1000) and TBR1 (Sigma, ab2261, 1:500) were diluted in blocking solution and incubated with tissue sections at 4 °C for 48 hours. Sections were then washed with 1X PBS (3 x 5 minutes), and incubated with the appropriate fluorescently labelled donkey- or goat-raised secondary antibodies (Invitrogen, 1:200) at room temperature for one hour. Sections were washed again with 1X PBS (3 x 5 minutes). We incubated sections with Hoechst solution in 1X PBS (Invitrogen, H3570, 1:1000) to stain cell nuclei at room temperature for five minutes. We then incubated sections in 0.01% Sudan black (Sigma Aldrich) solution at room temperature for five minutes. Sections were washed with 70% ethanol (1 x 30 seconds) and with 1X PBS (1 x 5 minutes), then mounted using Fluorescent Mounting Medium (Dako, S3023) without DAPI, and covered with 40 mm coverslips.

## Data availability

The raw and processed chimpanzee left frontal pole Visium spatial transcriptomics data were deposited in the Max Planck Institute Archive and are available on request (https://hdl.handle.net/1839/00f642f3-ac12-44e3-a3d3-f384884dc3e7).

## Code availability

All scripts used for the analysis are available on the project GitHub repository (https://github.com/galagoz/chimpbrain-st).

## Acknowledgements

We thank Zhiqiang Sha for his advice in spatial transcriptomics data analysis, Shaojie Ma and Andre M. M. Sousa for helping with chimpanzee dorsolateral prefrontal cortex single-nucleus RNA-sequencing data accession, and Else Eising and Barbara Molz for providing critical feedback on the manuscript. G.A., M.W., C.F., P.G., and S.E.F. are supported by the Max Planck Society (MPS). Brain tissue collection was supported by the inter-institutional funds of the President of the MPS, for the Hominoid Brain Connectomics (now Evolution of Brain Connectivity: EBC) Project. The funders had no role in study design, data collection and analysis, the decision to publish, or the preparation of the manuscript. S.E.F. is a member of the Center for Academic Research and Training in Anthropogeny (CARTA).

## Author contributions

G.A., M.W., and S.E.F. designed research; J.K. provided the *post mortem* tissue; T.G. and C.J. performed necropsy and tissue freezing; C.J. and M.M. performed tissue block cutting and sample shipment; M.W. and G.A. generated the data; G.A. analysed the data; G.A. wrote the initial draft of the paper; M.W., R.M.W., C.C., P.G., and S.E.F. supervised research; all co-authors provided critical feedback and commented on the manuscript.

