## Supplementary Materials for "Spatial and single-nucleus transcriptomic profile of a chimpanzee frontal pole"

**This PDF file includes:**

**Supplementary Figures 1-8**
**Supplementary Tables 1-4**

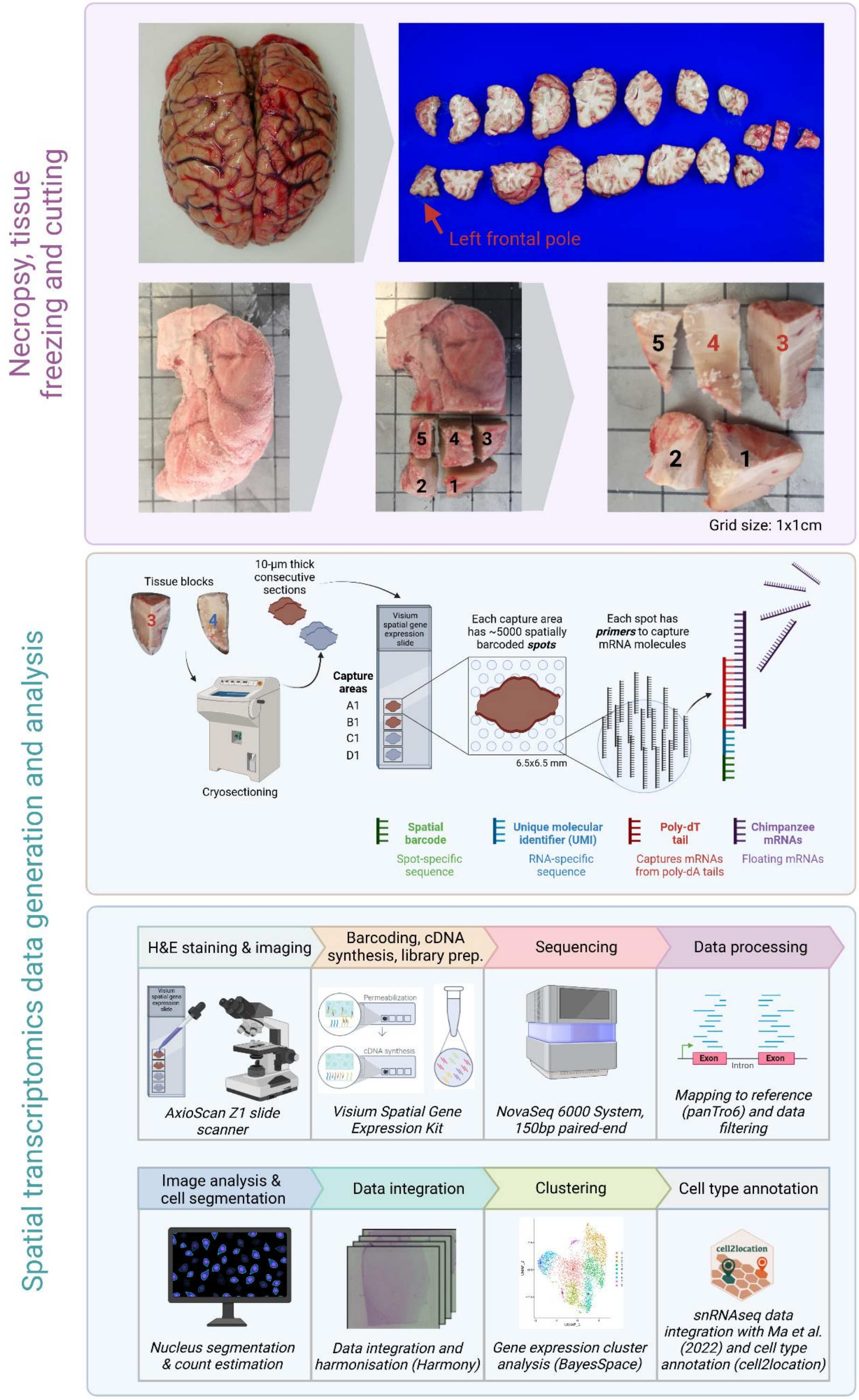


**Supplementary Figure 1:** Tissue extraction, freezing, cutting, spatial transcriptomics data generation and analysis workflow.

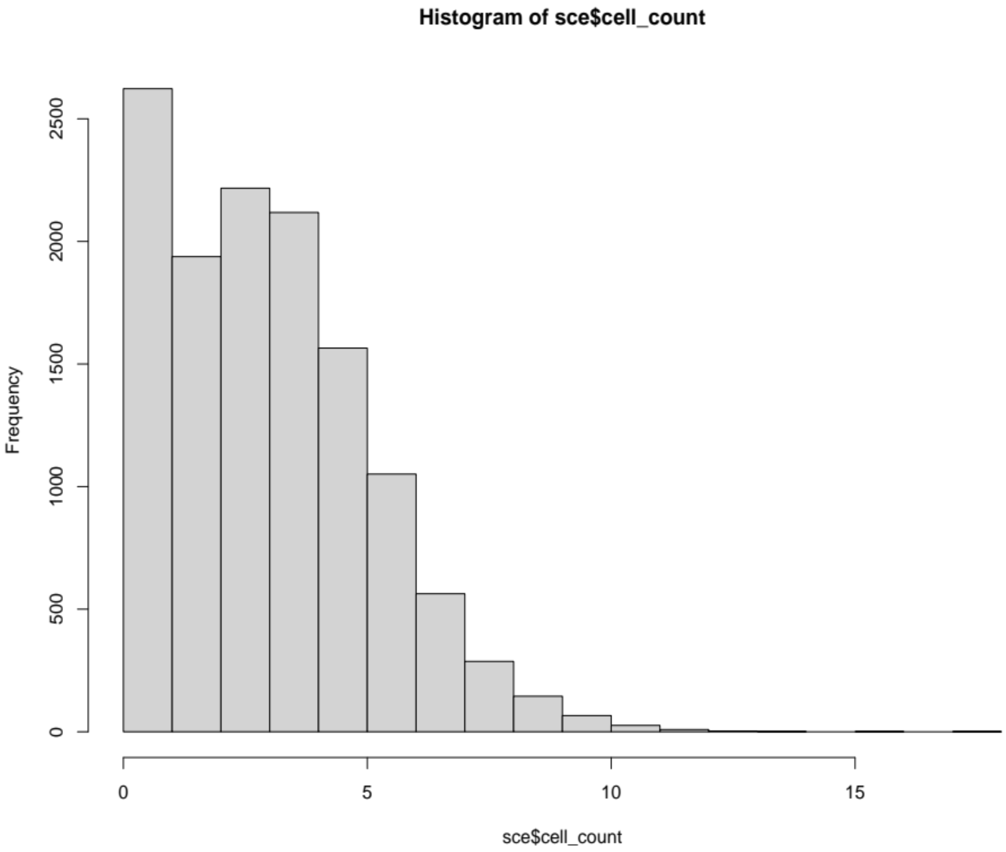


**Supplementary Figure 2:** Histogram of per-spot cell count estimates from across four tissue sections.

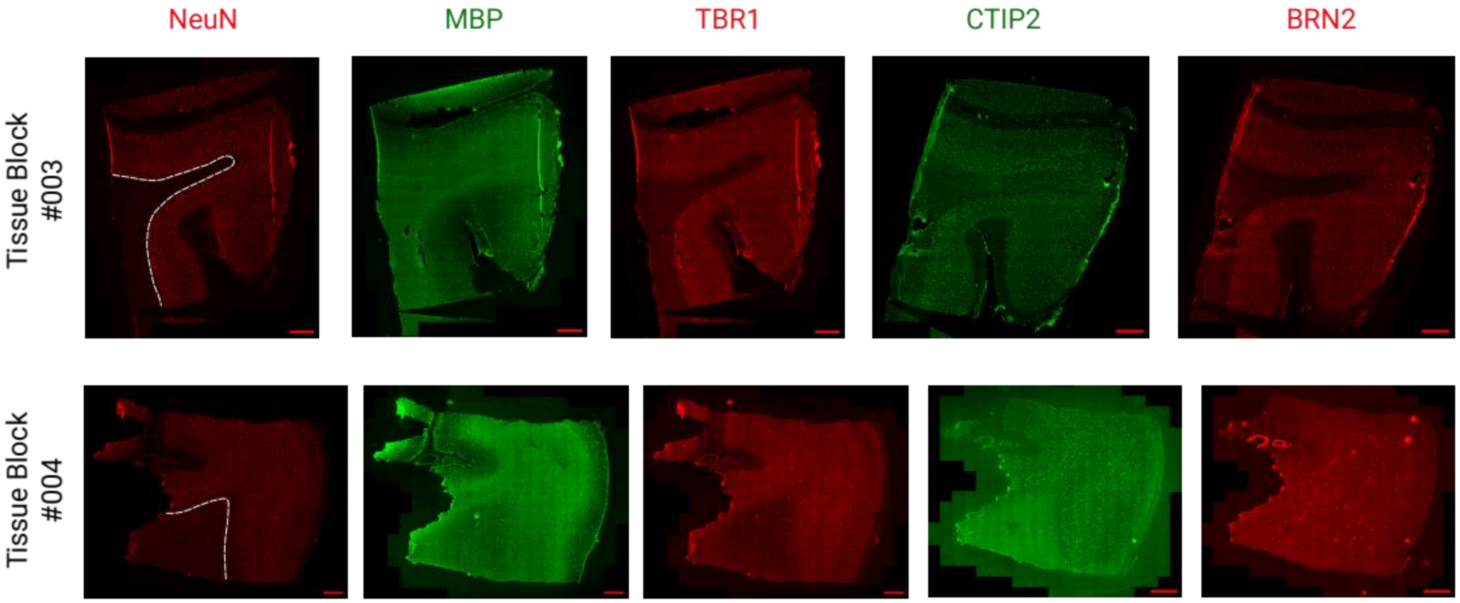


**Supplementary Figure 3:** Immunohistochemistry images of tissue sections that are adjacent to our Visium sections from tissue blocks #3 and #4. White-dotted lines indicate the white matter-grey matter border. Scale bar is 1 mm.

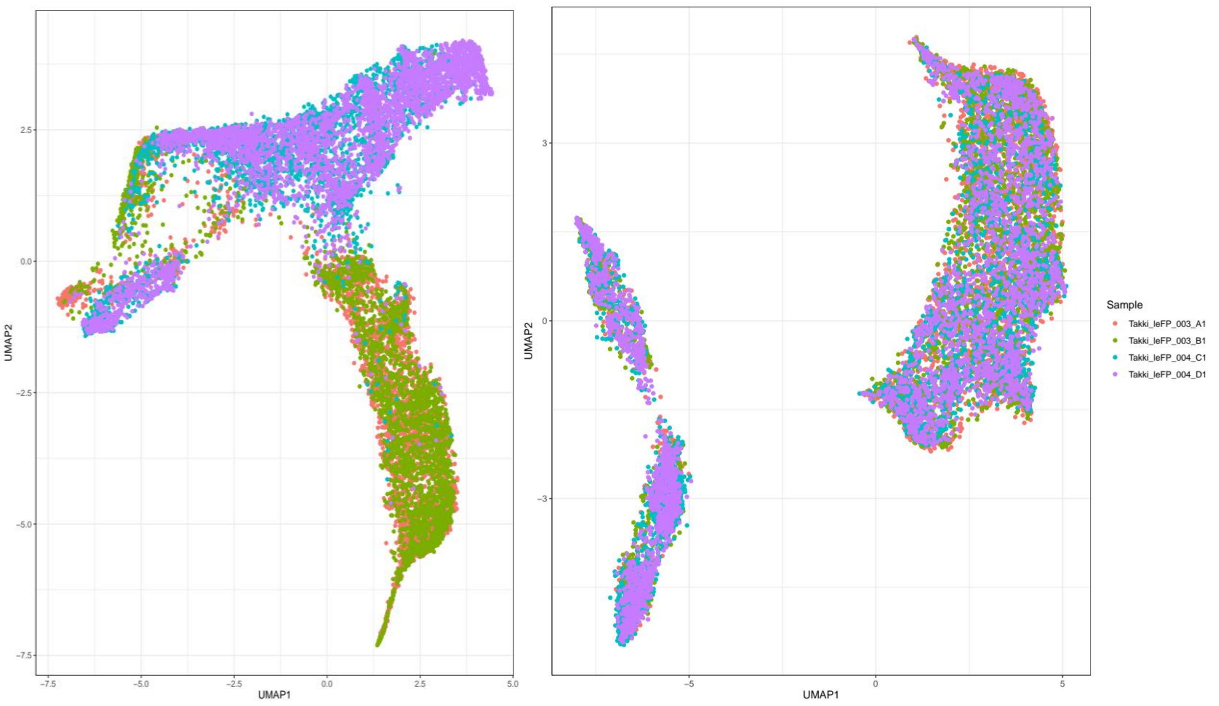


**Supplementary Figure 4:** Unsupervised clustering of Visium spots across four sections prior to (left) and after (right) batch effect correction by using Harmony.

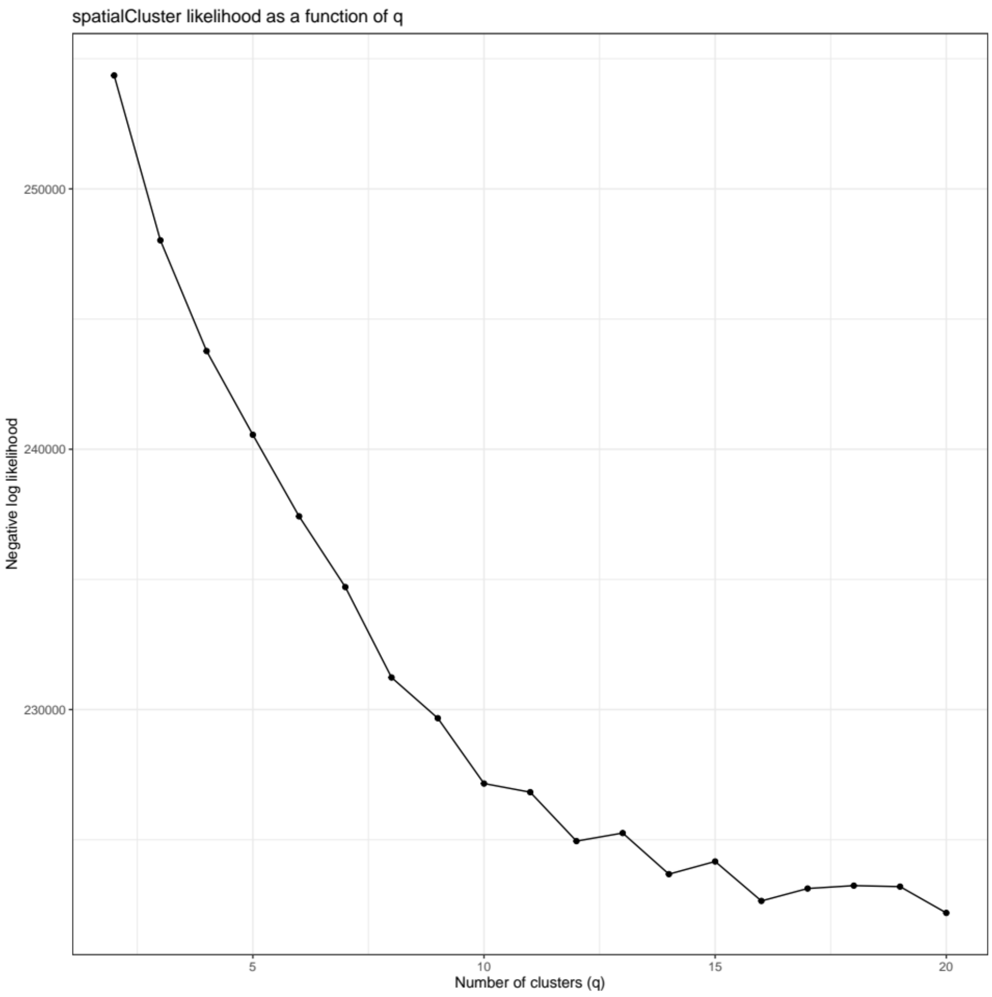


**Supplementary Figure 5:** The elbow plot showing negative log-likelihood estimates as a function of number of clusters (q).

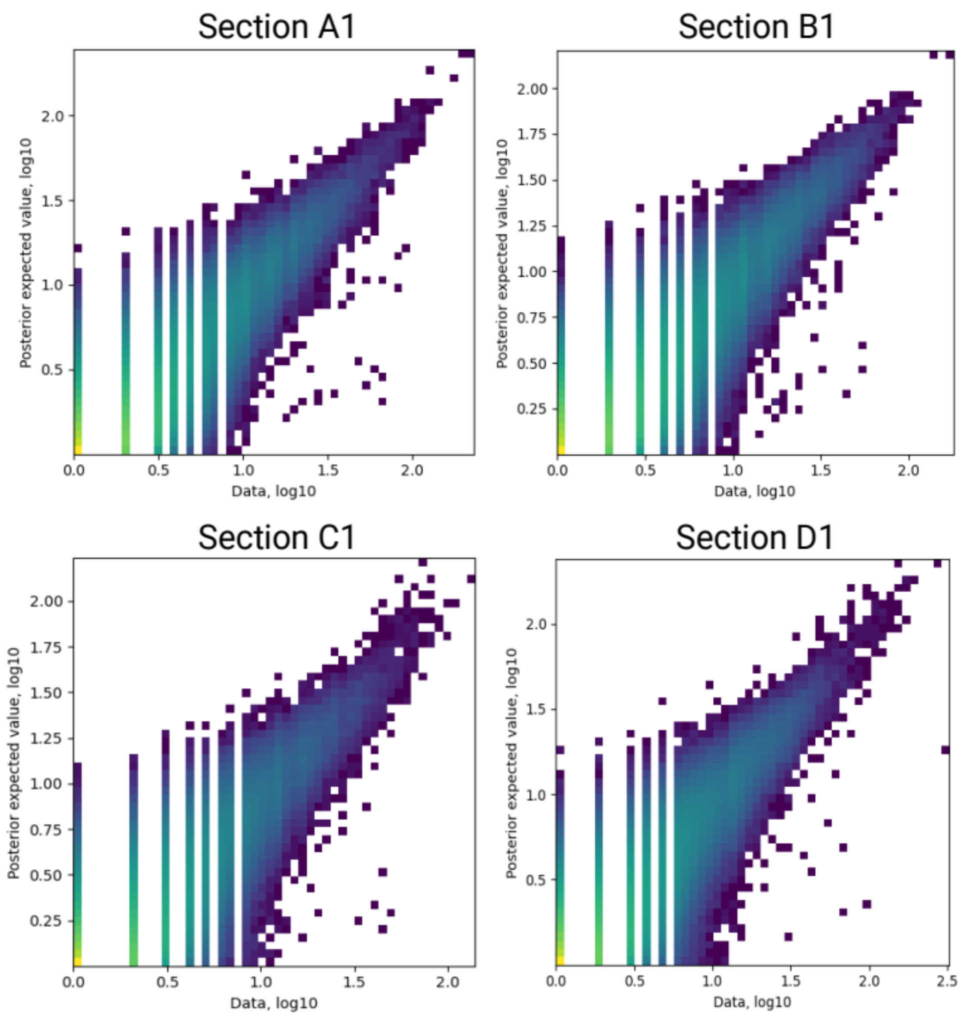


**Supplementary Figure 6:** cell2location model reconstruction accuracy 2D histograms for each section. Colour indicates 2D histogram count along x and y axes.

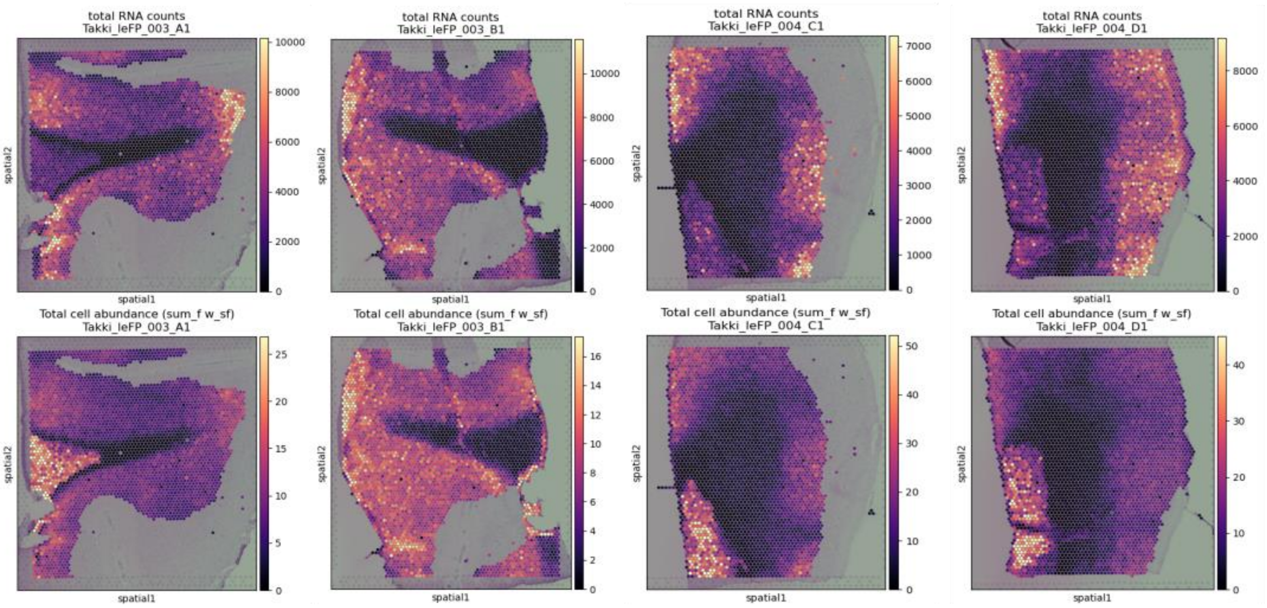


**Supplementary Figure 7:** Total RNA count and total cell abundance estimates per spot from cell2location quality control step.

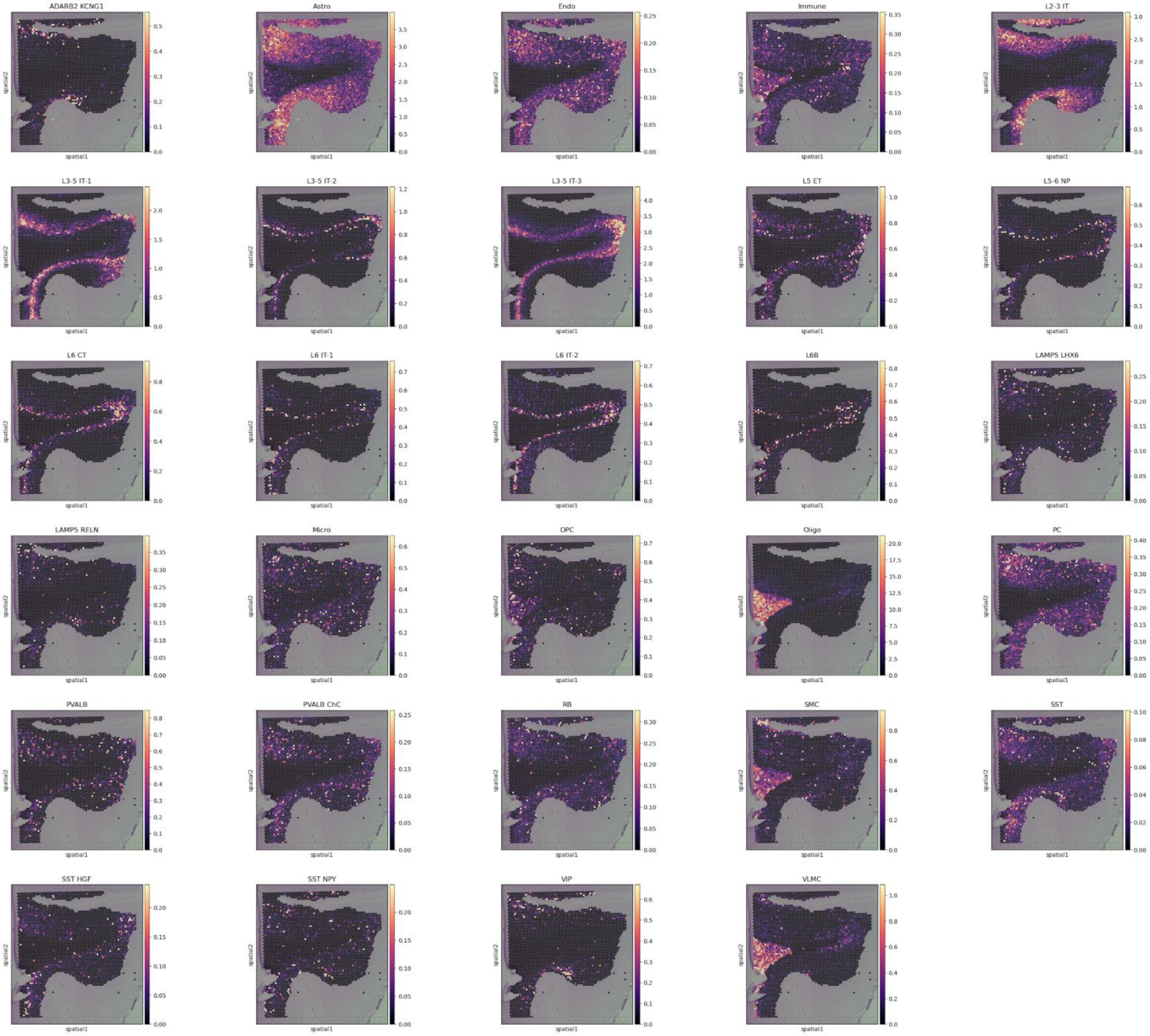
ADARB2 KCNG1
Astro
Endo
Immune
L2-3 IT
L3-5 IT-1
L3-5 IT-2
L3-5 IT-3
L5 ET
L5-6 NP
L6 CT
L6 IT-1
L6 IT-2
L6B
LAMP5 LHX6
LAMP5 RELN
Micro
OPC
Oligo
PC
PVALB
PVALB ChC
RB
SMC
SST
SST HGF
SST NPY
VIP
VLMC
spatial1
spatial2

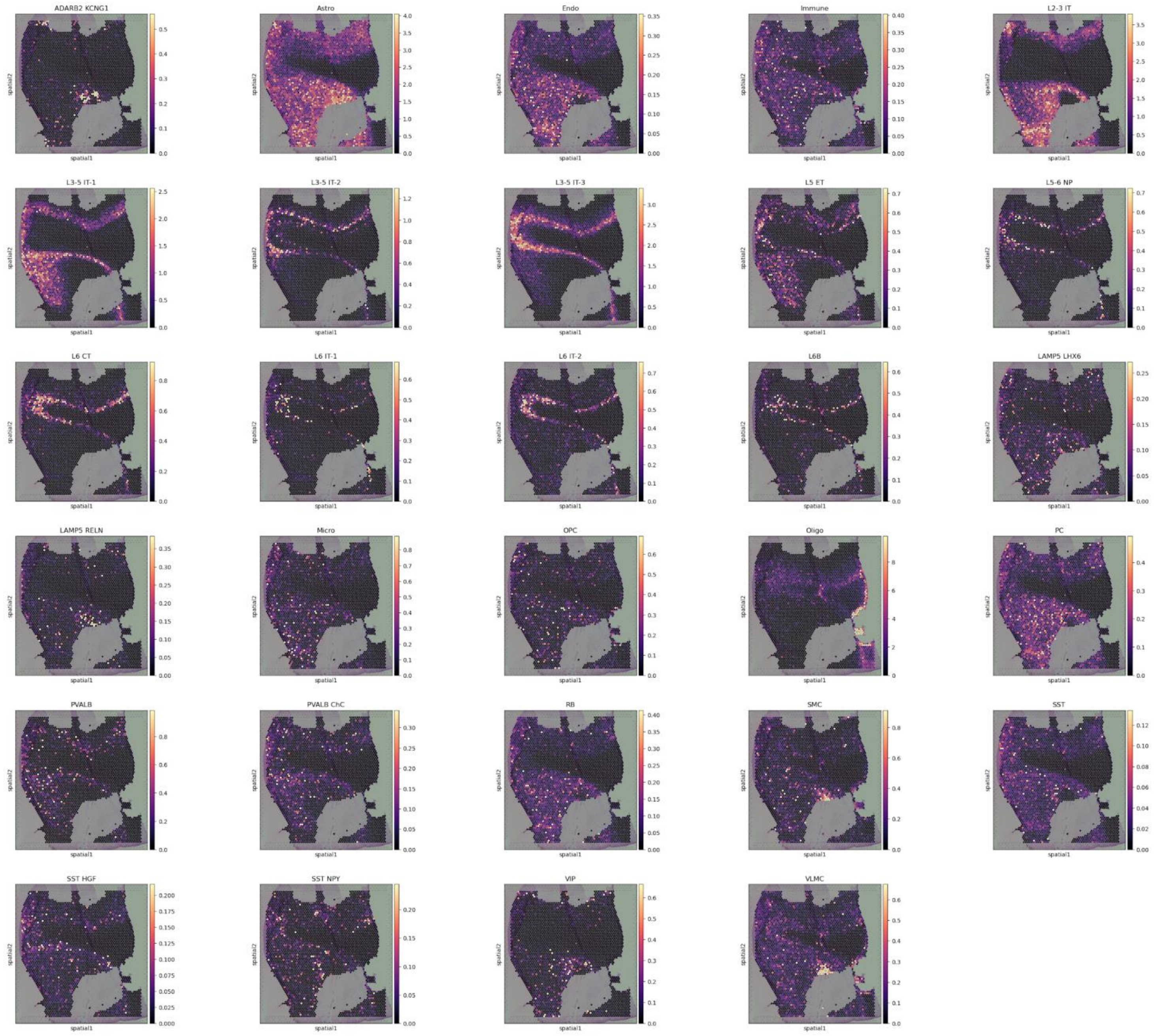
ADARB2 KCNG1
Astro
Endo
Immune
L2-3 IT
L3-5 IT-1
L3-5 IT-2
L3-5 IT-3
L5 ET
L5-6 NP
L6 CT
L6 IT-1
L6 IT-2
L6B
LAMP5 LHX6
LAMP5 RELN
Micro
OPC
Oligo
PC
PVALB
PVALB ChC
RB
SMC
SST
SST HGF
SST NPY
VIP
VLMC
spatial1
spatial2

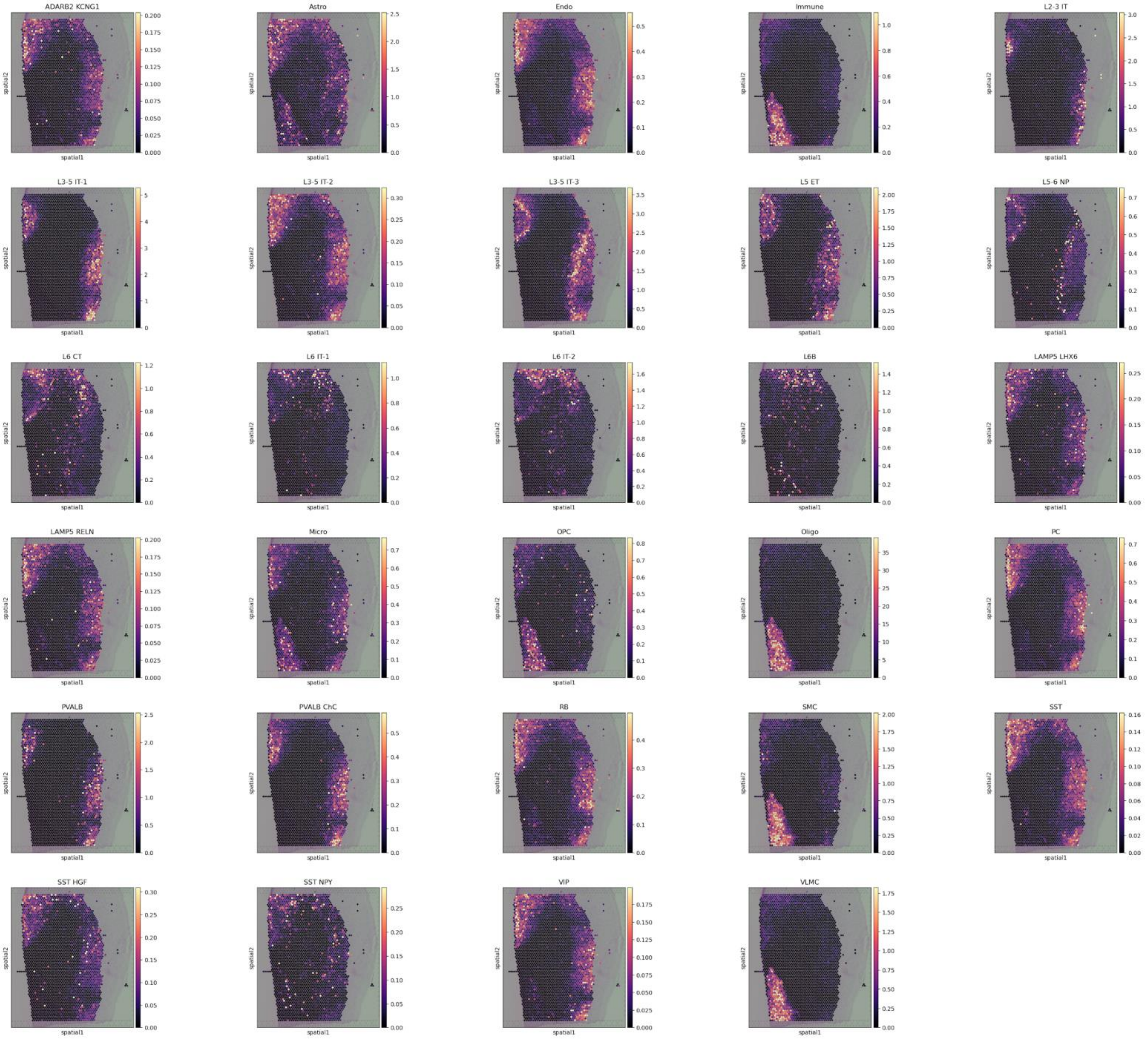
ADARB2 KCNG1
Astro
Endo
Immune
L2-3 IT
L3-5 IT-1
L3-5 IT-2
L3-5 IT-3
L5 ET
L5-6 NP
L6 CT
L6 IT-1
L6 IT-2
L6B
LAMP5 LHX6
LAMP5 RELN
Micro
OPC
Oligo
PC
PVALB
PVALB ChC
RB
SMC
SST
SST HGF
SST NPY
VIP
VLMC
spatial1
spatial2

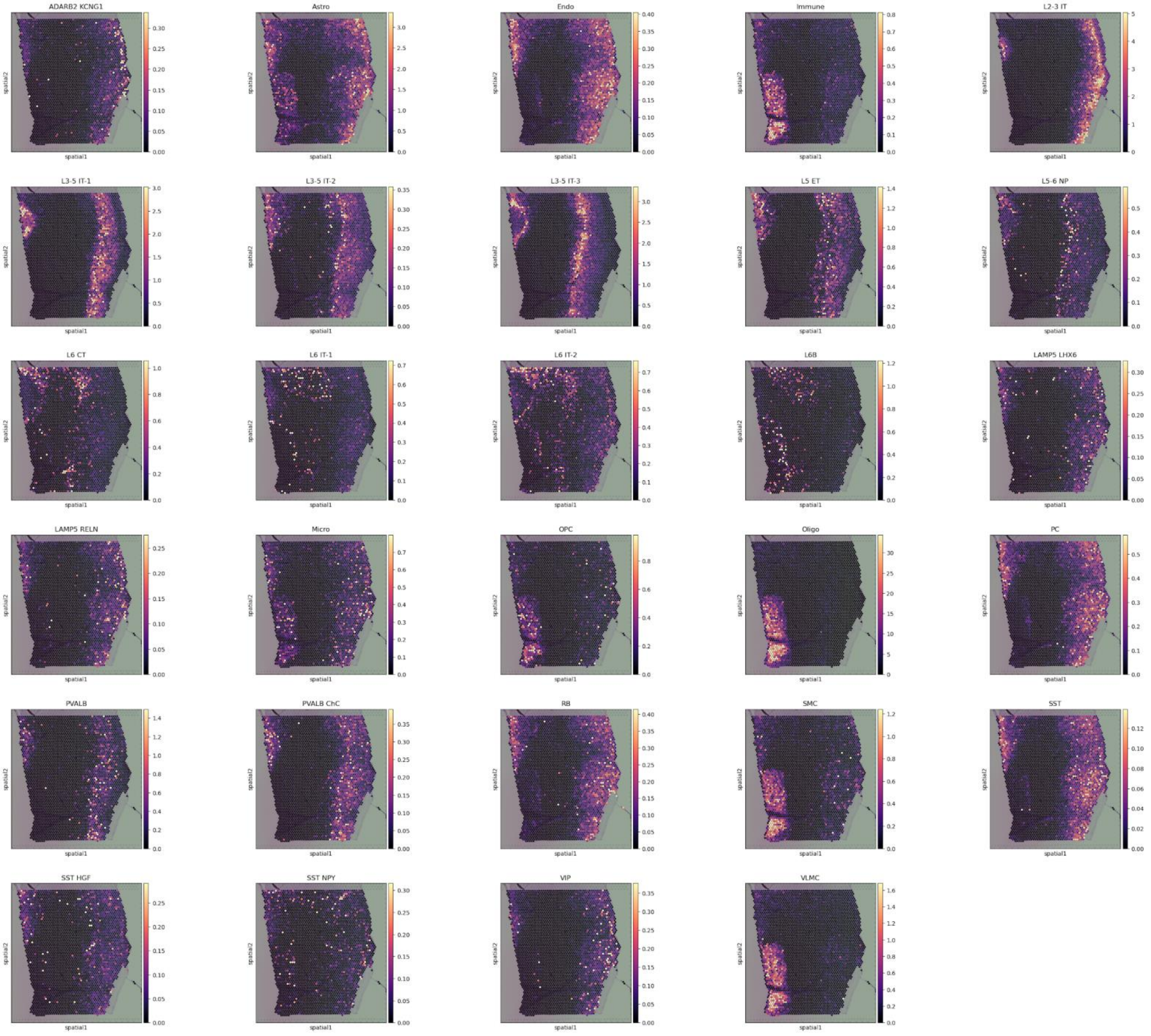


**Supplementary Figure 8:** Cell abundance estimates for 29 cell subclasses from Ma et al. (2022) per Visium sections A1, B1, C1, D1, respectively. Long forms of cell subclass abbreviations are described at Table S2.

| | | Nanodrop | | | Bioanalyser | |
|---|---|---|---|---|---|---|
| **Sample** | **Sample weight** | **RNA concentration (ng/ul)** | **260/280** | **260/230** | **RNA integrity number** | **rRNA Ratio (28s/18s)** |
| Takki_left_FP_002 | 10mg | 291,8 | 2,03 | 1,98 | 7,7 | 1,176322 |
| Takki_left_FP_005 | 30mg | 306 | 2,05 | 1,66 | 8,6 | 1,042234 |

**Supplementary Table 1:** Nanodrop and Bioanalyser RNA quantity and quality measurements from tissue blocks #2 and #5 (tissue blocks with corresponding IDs can be seen in Supplementary Figure 1).

| Capture Area | A1 | B1 | C1 | D1 |
|---|---|---|---|---|
| Tissue Block ID | Takki_leFP_003 | Takki_leFP_003 | Takki_leFP_004 | Takki_leFP_004 |
| Replicate | 1 | 2 | 1 | 2 |
| Position | 300 µm | 310 µm | 300 µm | 310 µm |
| Number_of_spots_under_tissue | 3.116 | 3.499 | 3.049 | 3.735 |
| Mean_reads_per_spot | 158.703 | 164.747 | 130.826 | 108.204 |
| Median_genes_per_spot | 2.122 | 2.545 | 1.157 | 1.691 |
| Number_of_reads | 494.518.529 | 576.448.275 | 398.887.795 | 404.140.278 |
| Valid_barcodes | 97,80% | 97,90% | 98,20% | 98,00% |
| Valid_UMIs | 99,90% | 99,90% | 99,90% | 100,00% |
| Sequencing_saturation | 88,40% | 89,40% | 93,80% | 91,50% |
| Q30_bases_in_barcode | 97,20% | 97,20% | 96,90% | 95,00% |
| Q30_bases_in_RNA_read | 90,20% | 90,10% | 92,10% | 90,60% |
| Q30_bases_in_UMI | 96,40% | 96,30% | 94,40% | 90,80% |
| Reads_mapped_to_genome | 88,10% | 84,80% | 97,10% | 96,70% |
| Reads_mapped_confidently_to_genome | 78,90% | 75,90% | 91,30% | 90,60% |
| Reads_mapped_confidently_to_intergenic_regions | 16,60% | 16,30% | 15,00% | 14,80% |
| Reads_mapped_confidently_to_intronic_regions | 4,10% | 4,10% | 3,60% | 3,40% |
| Reads_mapped_confidently_to_exonic_regions | 58,10% | 55,50% | 72,70% | 72,40% |
| Reads_mapped_confidently_to_transcriptome | 51,90% | 49,50% | 69,20% | 68,70% |
| Reads_mapped_antisense_to_gene | 0,70% | 0,60% | 0,60% | 0,60% |
| Fraction_reads_in_spots_under_tissue | 68,20% | 83,10% | 59,10% | 84,60% |
| Total_genes_detected | 16.288 | 16.585 | 15.437 | 16.096 |
| Median_UMI_counts_per_spot | 5.926 | 7.301 | 2.407 | 4.374 |
| Mean_cells_per_spot | 3,2576 | 3,1406 | 1,8872 | 2,3479 |

**Supplementary Table 2:** SpaceRanger QC metrics for each capture area on Visium spatial transcriptomic slide (slide no. V13J17-280).

| Study | Median sequencing depth | Mean UMI per spot | Mean genes per spot | Mean cell count per spot |
|---|---|---|---|---|
| This study | $283\times10^6$ | 5002 | 1879 | 2,7 |
| Wong and Sha et al. (preprint, 2024) | $469\times10^6$ | 6785 | 2783 | NA |
| Maynard et al. (2021) | $291\times10^6$ | 3462 | 1734 | 3,3 |
| Chen et al. (2020) | NA | 8638 | 3340 | NA |

**Supplementary Table 3:** Comparison of various RNA-sequencing data quality measures between the current study and human brain spatial transcriptomics studies from literature.

| Class | Subclass | Description |
|---|---|---|
| Excitatory (glutamatergic) neuron | L2-3 IT | Layer 2-3 intratelencephalic |
| Excitatory (glutamatergic) neuron | L3-5 IT-1 | Layer 3-5 intratelencephalic Type 1 |
| Excitatory (glutamatergic) neuron | L3-5 IT-2 | Layer 3-5 intratelencephalic Type 2 |
| Excitatory (glutamatergic) neuron | L3-5 IT-3 | Layer 3-5 intratelencephalic Type 3 |
| Excitatory (glutamatergic) neuron | L5 ET | Layer 5 extratelencephalic |
| Excitatory (glutamatergic) neuron | L5-6 NP | Layer 5-6 near-projecting |
| Excitatory (glutamatergic) neuron | L6 IT-1 | Layer 6 intratelencephalic Type 1 |
| Excitatory (glutamatergic) neuron | L6 IT-2 | Layer 6 intratelencephalic Type 2 |
| Excitatory (glutamatergic) neuron | L6 CT | Layer 6 corticothalamic |
| Excitatory (glutamatergic) neuron | L6B | Layer 6 Type B |
| Inhibitory (GABAergic) neuron | LAMP5 LHX6 | *LAMP5*- and *LHX6*-expressing cells |
| Inhibitory (GABAergic) neuron | LAMP5 RELN | *LAMP5*- and/or *RELN*-expressing cells |
| Inhibitory (GABAergic) neuron | VIP | *VIP*-expressing cells |
| Inhibitory (GABAergic) neuron | ADARB2 KCNG1 | *ADARB2*- or *KCNG1*-expressing cells |
| Inhibitory (GABAergic) neuron | SST | *SST*-expressing cell |
| Inhibitory (GABAergic) neuron | SST NPY | *SST*- and *NPY*-expressing cells |
| Inhibitory (GABAergic) neuron | SST HGF | *SST*- and *HGF*-expressing cells |
| Inhibitory (GABAergic) neuron | PVALB | *PVALB*-expressing cells |
| Inhibitory (GABAergic) neuron | PVALB ChC | *PVALB*- and *ChC*-expressing cells |
| Glia | Astro | Astrocytes |
| Glia | OPC | Oligodendrocyte precursor cells |
| Glia | Oligo | Oligodendrocytes |
| Glia | Micro | Microglie |
| Non-neuronal cell | Immune | Immune cells (macrophage, myeloid, B & T cells) |
| Non-neuronal cell | Endo | Endothelial cells |
| Non-neuronal cell | RB | Red blood lineage cells |
| Non-neuronal cell | PC | Pericytes |
| Non-neuronal cell | SMC | Smooth muscle cells |
| Non-neuronal cell | VLMC | Vascular leptomeningeal cells |

**Supplementary Table 4:** Abbreviations and descriptions of 29 cell subclasses identified by from Ma et al. (2022).
